# Molecular cloning and recombinant protein expression of 9-Lipoxygenase gene from *Solanum tuberosum* in *Pichia pastoris* yeast cells

**DOI:** 10.1101/2022.05.18.492528

**Authors:** Alex Olengo, Ann Nanteza, Elvio Henrique Benatto Perino, Leif-Alexander Garbe, Sylvestre Marillonnet, Ramona Grützner, Margaret Saimo-Kahwa, Fabien Schultz, Matthias Koch

**Author notes:** Corresponding author: Alex Olengo.

## Abstract

Synthetic fungicides have effectively controlled plant pathogenic fungi but their continued use has been detrimental to natural biological systems, and sometimes resulted into development of fungal resistance. They also have undesirable effects on non-target organisms and foster environmental and human health concerns thus new biodegradable alternatives have to be investigated. Lipoxygenases (LOX) are ubiquitous non-heme Iron containing dioxygenases that catalyze the addition of molecular oxygen to Polyunsaturated Fatty Acids (PUFAs) such as Linoleic acid to form Oxylipins that possess anti-microbial activity.The aim of this study was to generate a recombinant 9-Lipoxygenase protein for chemo enzymatic synthesis of Oxylipin based biodegradable fungicides. Golden gate assembly, a molecular cloning method that allows assembly of many DNA fragments into a complete piece using Type II s restriction enzymes and T4 DNA ligase was used to clone the complete 9-LOX gene into the expression plasmid vector pPICZαB. Protein expression in *Pichia pastoris* yeast cells was induced by addition of absolute methanol every after 24h for up to four days. Analysis of protein expression from cell lysates was achieved by SDS-PAGE and Western blotting probed with anti-histidine antibody which showed putative protein bands of 97kDa representing recombinant 9-LOX protein. It is recommended that optimization studies on the yeast kex2 convertase and the α- secretion factor can be done to enable secretion of recombinant *Solanum tuberosum* 9-LOX protein since the protein in this study was recovered from cell lysates.

## Introduction

Lipoxygenases (LOXs) belong to a family of monomeric proteins that catalyze oxidation of polyunsaturated fatty acids (PUFA) (linoleic, linolenic, and arachidonic acid) to form hydroperoxides (Lončarić *et al*., 2021). The LOXs are ubiquitous in animal, plant, and fungi kingdoms, as well as in cyanobacteria (Newcomer & Brash, 2015). During insertion of molecular oxygen into the polyunsaturated fatty acids, the oxygen can be added to both sides of the pentadiene system a term known as Regio specificity (Conconi *et al*., 1996). According to Brash (1999), addition of molecular oxygen to the PUFA can either happen at carbon position 9 or 13 and this yields two products (9 or 13-hydroperoxy fatty acids) respectively. The unsaturated fatty acid hydroperoxides produced as a result of LOX activity are further metabolized into other lipid mediators for example in plants, the mediators yielded include volatile aldehydes and jasmonates (Feussner & Wasternack, 2002). These lipid mediators have been shown in several studies to confer antimicrobial action to fungal and bacterial attack for example in tobacco following infection with *Phytophthora parasitica var. nicotianae* and in potato infected by *Phytophthora infestans* (Polkowska-Kowalczyk *et al*., 2008). More so, the accumulation of the 9-LOX-derived oxylipins from *Solanum tuberosum* has been shown to be rapid and intense in stimulation of plant defence gene expression, modulation of cell death and/or showing antimicrobial activities hence, they are considered to be potential defence compounds (Polkowska-Kowalczyk *et al*., 2008). Synthetic fungicides have a problem of bioaccumulation and non-biodegradability thus there is need to deploy techniques for production of natural fungicides such as oxylipins that confer antifungal properties to field crops with no bioaccumulation. Therefore, the Golden gate cloning strategy was used in this study to assemble the complete 9-LOX gene which was subsequently cloned into pPICZαB expression vector.

After cloning, *Pichia pastoris* X-33 expression cell system was utilized to express the recombinant protein of interest. Golden Gate cloning is a method used to assemble multiple DNA fragments in a defined linear order into a recipient vector using a one-pot assembly procedure. The principle of golden gate cloning is based on the use of a Type IIs restriction enzyme for digestion of the DNA fragments and vector (Marillonnet & Grützner, 2020). *Pichia pastoris* is a methylotrophic yeast implying that it can grow with only methanol as a source of energy. The gene that codes for the desired protein is mainly achieved by inducing AOX1 promoter through addition of absolute methanol (Koutz *et al*., 1989). The yeast system has short expression time, cost effective, and cotranslational and posttranslational processing (Karbalaei, Rezaee, & Farsiani, 2020). Studies have recently indicated that the *Pichia* expression system is unique in the expression of membrane proteins ranging from calcium and potassium channels, nitrate and phosphate transporter, and histamine H1 receptor (Karbalaei *et al*., 2020a). This study has been able to generate a cell line that can be scaled up for production of more recombinant 9-LOX protein for chemo enzymatic synthesis of oxylipin based fungicides. This could tackle the challenge of nonbiodegradable fungicides used by farmers in crop fields hence preserving the ecosystem integrity.

## Materials and Methods

### Sources of materials used in the study

The recombinant PET-28a-9-LOX plasmid vector (Royo *et al*., 1996) used as a template for 9-LOX amplification in this study was donated by Professor Sabine Rosahl of the Leibniz Institute of Plant Biochemistry (Halle, Germany). The intermediary vectors used in the cloning (pAGM35763) and plasmid pICH41308 (PMC3041749) used for *Lac Z* amplification work were generously provided by Sylvestre Marillonnet of the Leibniz Institute of Plant Biochemistry, Germany. Primers were designed using Vector NTI Version 11.5.5 (Thermo Fisher Scientific) and purchased from Fermentas GmbH, Germany. The PCR kit used in the study was also purchased from Fermentas GmbH. The expression kit containing pPICZαB plasmid was purchased from Thermo Fisher Scientific, Germany. Primary and secondary antibodies used in the Western blot analysis were purchased from Abcam, England. The restriction enzymes (*BsaI, LguI*, and *PmeI*) were purchased from New England Biolabs whereas *EcoRI* and *X-bal* were purchased from Thermo Fisher Scientific. Ligase enzyme and buffer were purchased from Promega. NucleoSpin^®^ Plasmid Easy Pure kit was purchased from Macherey Nagel.

### PCR amplification of three defined 9-LOX gene fragments from PET-28a-9-LOX DNA

The three defined 9-LOX gene fragment regions selected by the Vector NTI software were amplified by PCR using the 3 pairs of primers (Table 1) which added *Bsa I* sites flanking both ends of each of the three 9-LOX amplicons for restriction-ligation cloning into intermediary vector *pAGM35763* using *BsaI* Type II restriction enzyme. The PCR was performed using KOD Hot start DNA polymerase (Millipore Sigma) since it has a low error rate unlike Taq polymerase (Engler and Marillonnet 2013). For this purpose therefore, three PCR tubes were used and reaction was set up using PCR conditions as described by Engler and Marillonnet (2013). Briefly, each of the three PCRs was set up in a volume of 50 μl containing 50ng of PET-28a-9-LOX DNA, 200μM of each of dNTPs (Fermentas), 1x of Advantage^®^ 2 Buffer (Clonetech), 1.5mM MgSO4, 0.3μM of each of the sense and antisense primers, 1.25 U of KOD hot start DNA polymerase (Millipore Sigma) and nuclease free water. The PCR was performed in a BioRad thermocycler (Biorad-Hercules, California). The PCR program was set up with an initial denaturation at 95°C for 2 min, followed by 35 cycles of denaturation at 95°C for 20 s, primer annealing at 58°C for 10 s and extension at 70°C for 2 min. This was followed by a final extension step at 70°C for 5 min and holding at 10°C. From the amplified products, 5μl of each of the three 9-LOX amplicons were mixed with 1μl of 6X gel loading dye (Thermo-Fisher Scientific), loaded on a 1% agarose gel and electrophoresed against a 1kb DNA ladder (RTU-Thermo Scientific) at 125V in 1X Tris-Acetic acid-EDTA (TAE) buffer containing 0.5*μ*g/ml ethidium bromide for 35 minutes and visualized using the UV gel documentation system -Wagtec, United Kingdom.

**Table 1:**
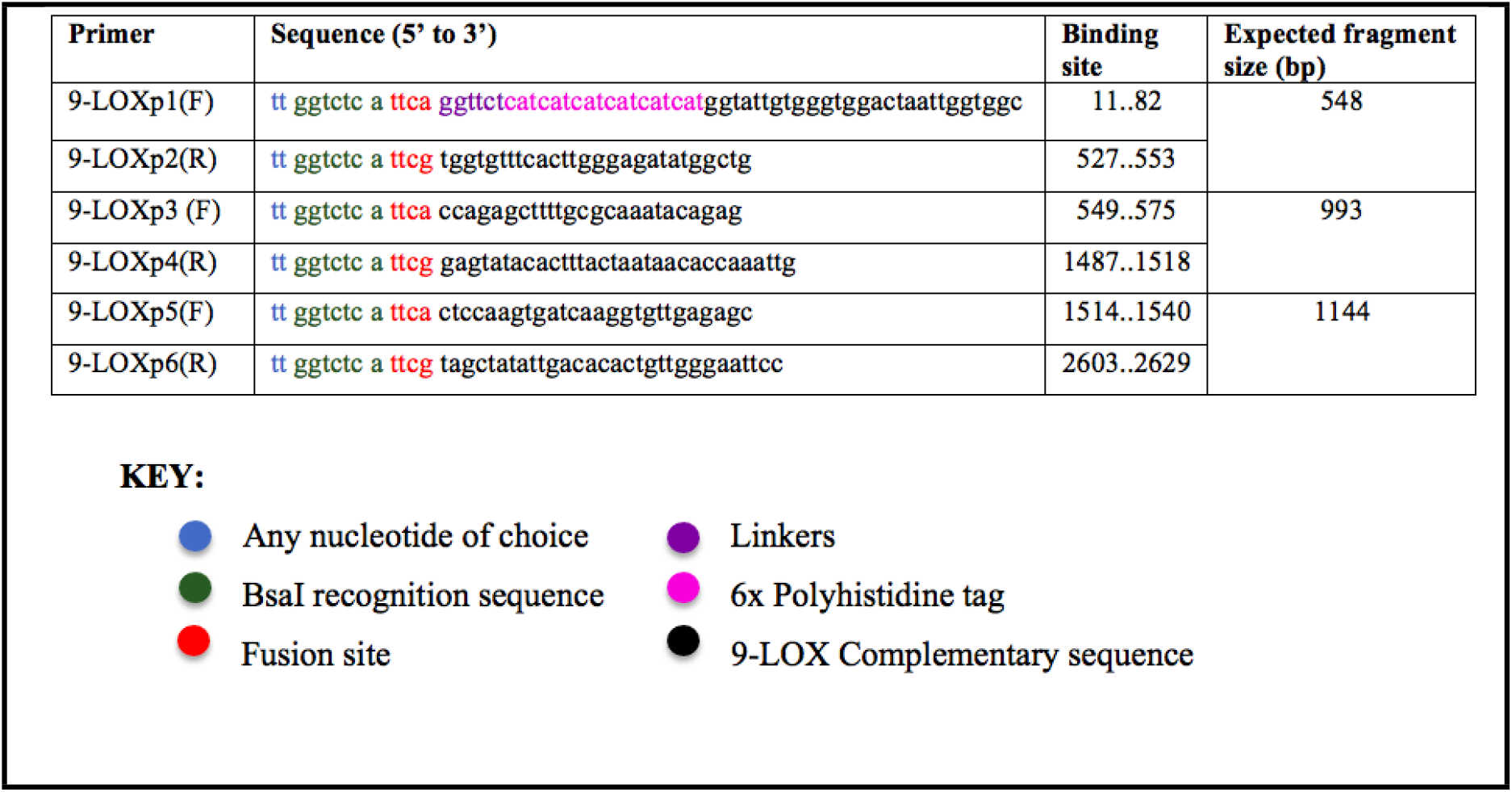
Primer pairs used for amplification of three 9-LOX gene fragments from *Solamum tuberosum*

From the remaining 45μl of each of the 3 PCR amplicons, 10μl was loaded on a 1% agarose gel and run at same conditions as for the initial. The DNA bands were then excised from the gel and purified of the remaining primers, potential dimers, and also remaining polymerase enzyme by use of Nucleospin^®^ Extract II Kit (Clontech Laboratories) following Manufacturer’s protocol. The concentration of the purified DNA was measured using Qubit^®^ 3.0 Fluorometer (Invitrogen) following manufacturer’s protocol.

### Cloning of three 9-LOX PCR fragments individually into the intermediary plasmid vector pAGM35763

The intermediary vector *pAGM35763* used for this cloning step had compatible *BsaI* sites to those of the three 9-LOX amplicons. The vector was used in a restriction-ligation reaction to generate intermediary recombinant vectors *pAGM52925* containing a 548bp fragment, *pAGM52901* containing a 993bp fragment and *pAGM52914* containing a 1144bp fragment (Figure 1).

**Figure 1:**
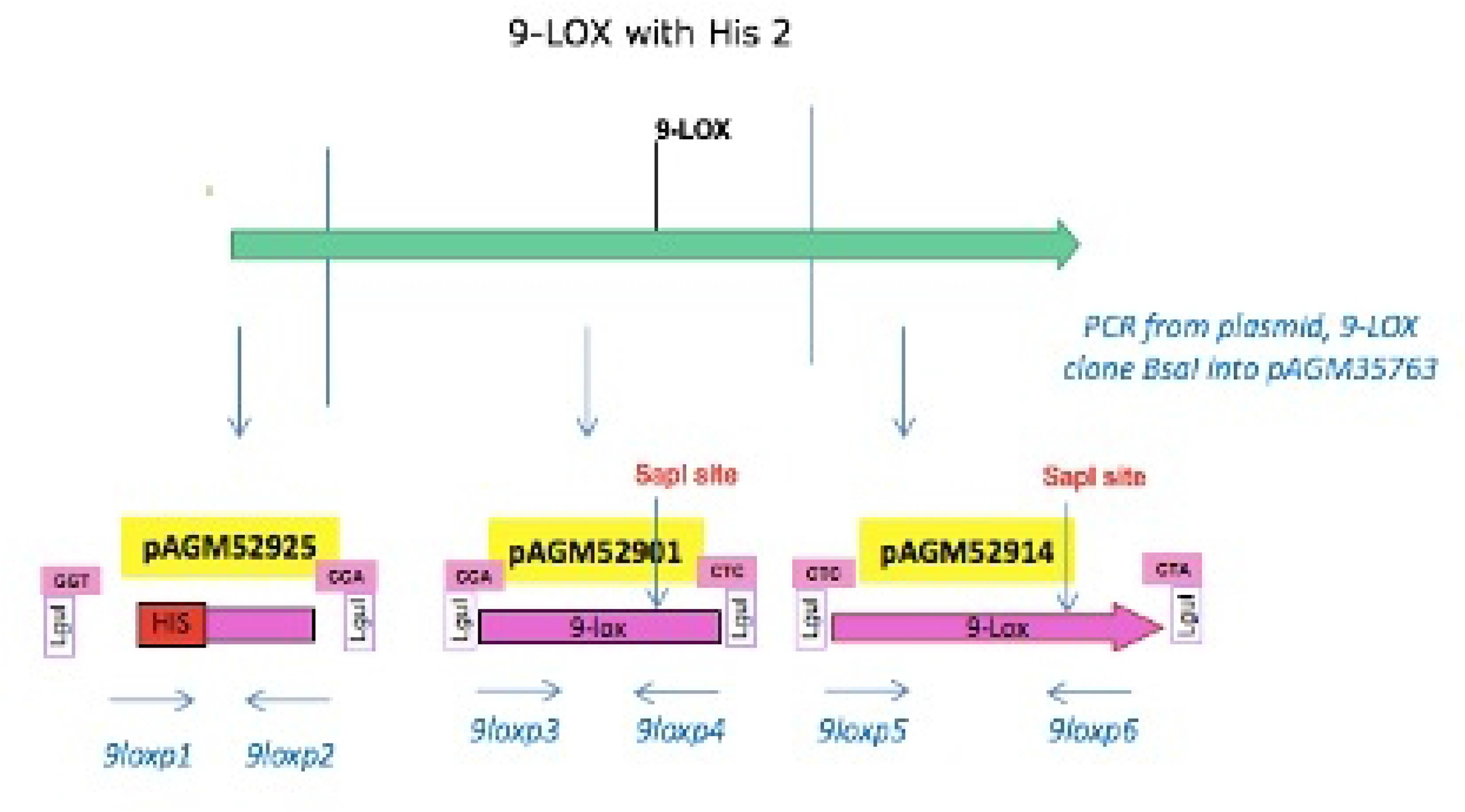
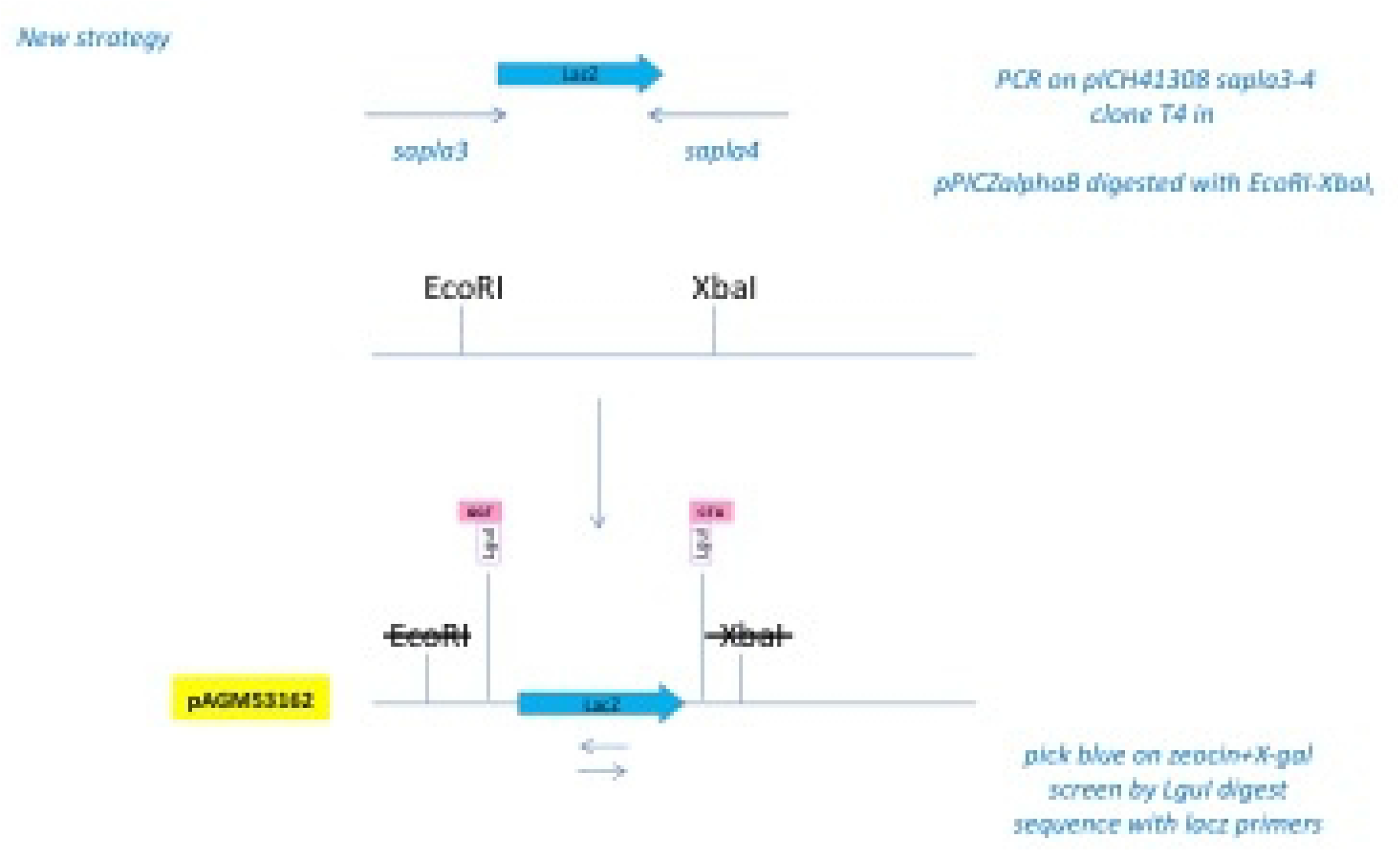
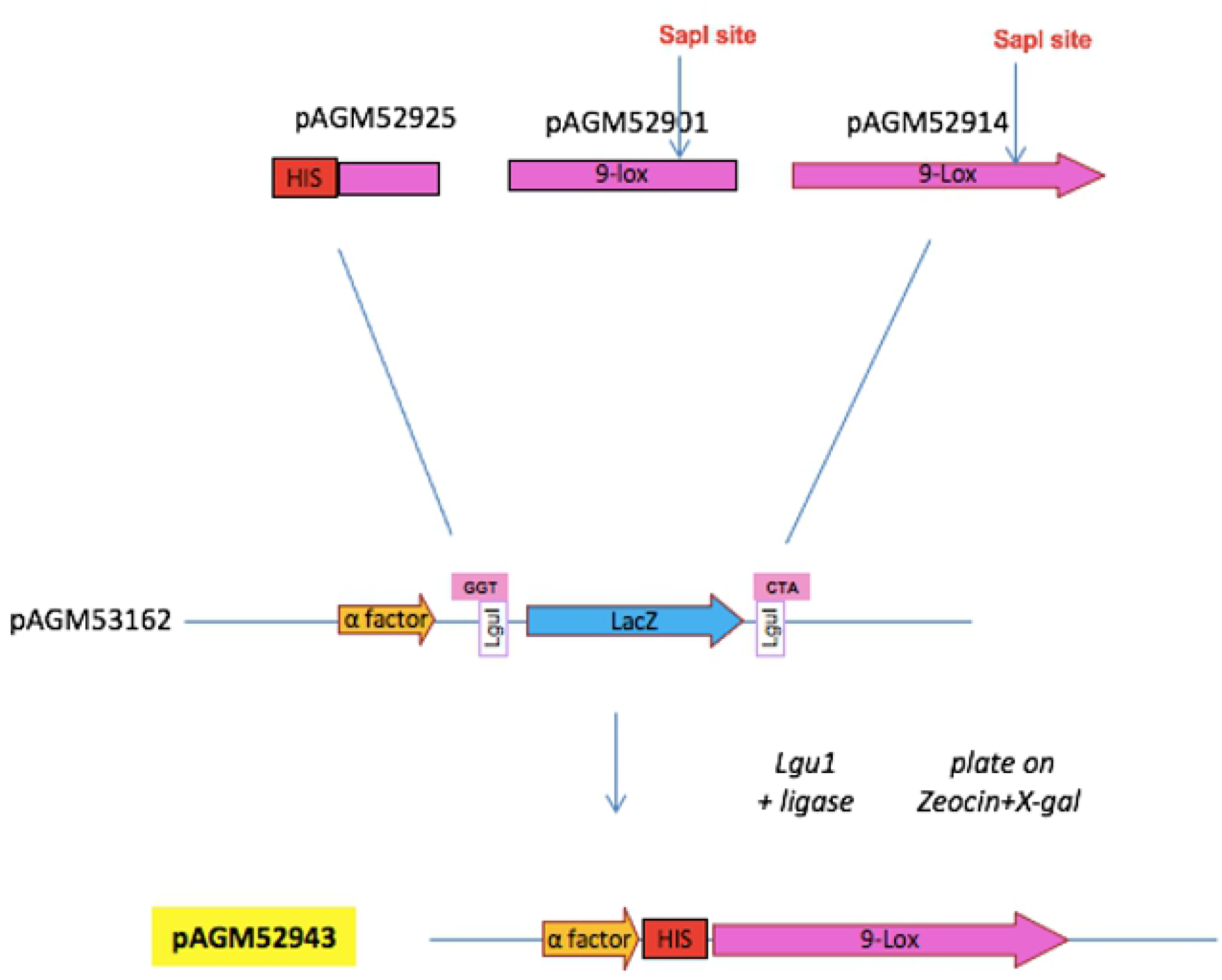
Schematic representation of the cloning steps. a) Amplified 9-LOX regions that were cloned into pAGM35763 Vector. b) Schematic representation of T4 cloning of LacZ amplicon into pPICZαB plasmid vector. c) Schematic representation of restriction-ligation cloning of 9-LOX fragments into pPICZαB expression vector

Briefly, in a 15μl restriction-ligation reaction for each of the 9-LOX amplicons and intermediary vector *pAGM35763*, the reaction was set up using a protocol adopted from Engler & Marillonnet (2013) as follows:50ng of vector, 50ng 9-LOX amplicon, 1x ligation buffer (Promega), 3U of *BsaI* enzyme (New England Biolabs), 1U T4 DNA ligase (Promega) and nuclease-free water. The reaction mixtures in the 3 PCR tubes were incubated at 37°C for 4 hours for initial restriction step of *BsaI* enzyme and ligase activity, followed by 50°C for 5min for a second restriction step of *BsaI*, then 80°C for 5min to inactivate the remaining enzymes. The reaction mixtures were held at 10°C until the PCR tubes were removed from the thermocycler (C1000 Touch, BioRad). After, freshly prepared chemically competent *E*.*coli* DH10B cells (100μl) were transformed with 10μl of each of the restriction-ligation products. Using a sterile wire loop, the transformants from each of the three tubes were spread on three separate LB agar plates and incubated overnight at 37°C in an oven incubator (Memmert IN110 PLUS, USA). White and blue colonies were observed but only 5 white colonies from each of the three LB plates were picked and individually inoculated in five 50ml falcon tubes containing 5 ml of LB medium with 25μg/ml of spectinomycin then grown overnight at 37°C. Plasmid DNA was then extracted from the 15 falcon tubes (5 tubes for each plasmid) using the NucleoSpin^®^ Plasmid Easy Pure kit from Macherey Nagel following the manufacturer’s instructions. After, concentration of the 15 plasmid DNA samples (5 for each recombinant intermediary vector) were done using Qubit^®^ 3.0 Fluorometer-Invitrogen. All the 15 plasmid samples were also sent to Eurofins Genomics (MWG) for sequencing to check whether the 15 plasmid samples were in the correct reading frame and had no mutations. One representative plasmid DNA sample from each of the recombinant vectors (*pAGM52925, pAGM52901* and *pAGM52914*) was selected for cloning into the destination vector (pPICZαB) using golden gate assembly procedure (Engler & Marillonnet, 2013).

### Preparation of destination vector for golden gate assembly procedure

#### LacZ fragment amplification and digestion of destination plasmid vector pPICZαB

The vector was digested with *EcoRI* and *Xbal* restriction enzymes (Figure 1b) then cloned with a *Lac Z* amplicon that introduced *LguI* restriction sites into the vector. The *Lgul* sites were used for complementary ligation of the 9-LOX fragments in the final cloning step into the destination vector (pPICZαB). The Lac Z gene was used for blue-white selection of colonies using x-gal.

The plasmid vector *pICH41308* was used as a template for *LacZ* gene amplification using sapla 3 forward primer (5’gaggctgaagctgcaggaattggtt**<u>gaagagc</u>**gcagctggcacgacaggtttc3’) and sapla 4 reverse primer (5’gatcctcttctgagatgagtttttgttctagt**<u>gaagagc</u>**gtcacagcttgtctgtaagcgg3’) that introduced *Lgu1* recognition sites (bold and underlined) compatible with the LguI sites in the intermediary vector used in subcloning of 9-LOX fragments. The *LacZ* amplification was performed using PCR conditions as described by Engler & Marillonnet (2013). A reaction volume of 50μl was set up as follows: 50ng of plasmid vector *pICH41308*, 200μM of each of dNTPs (Fermentas), 1.5mM MgSO4,1X of Advantage^®^ 2 Buffer (Fermentas), 0.3μM of each of the forward and reverse primers, 1.5 U of KOD hot start DNA polymerase (Fermentas) and nuclease-free water. The PCR was performed in a BioRad thermocycler (Biorad-Hercules, California). The PCR program was set up with initial denaturation at 95°C for 2min, followed by 35 cycles of denaturation at 95°C for 20s, primer annealing at 58°C for 10s and extension at 70°C for 2min. This was followed by a final extension step of 70°C for 5min and holding at 10°C. From the amplified product, 5μl was mixed with 6X gel loading dye (Thermo-Fisher Scientific), loaded on a 1% agarose gel and run alongside a 1kb DNA ladder (RTU-Thermo Scientific) at 125V in 1X Tris-Acetic acid-EDTA (TAE) buffer that contained 0.5*μ*g/ml ethidium bromide for 35 minutes. The ethidium bromide stained gel was visualized using the UV gel documentation system Wagtec, United Kingdom. A 649bp fragment which was the expected band size was detected (Figure 3). From the remaining 45μl of the amplified product, 10μl was loaded on a 1% agarose gel and run as above. The DNA band was cut out and purified using the Nucleospin^®^ Extract II Kit (Clontech Laboratories) protocol. After, DNA concentration was determined using Qubit^®^ 3.0 Fluorometer (Invitrogen) following manufacturer’s protocol.

#### T4 cloning of LacZ amplicon into digested pPICZαB plasmid vector

Basing on a protocol adopted from Jeong *et al* (2012), *EcoRI* and *XbaI* restriction digested pPICZαB vector and the purified *LacZ* PCR product were mixed together in a 20μl reaction volume containing: 100ng of digested pPICZαB vector, 40ng of the *LacZ*, 1X ligase buffer (Promega), 0.6 U of T4 DNA polymerase (Thermo Fisher Scientific) and nuclease free water. The contents in the PCR tube were incubated at 25°C for 5 min and then put on ice for 1 min to stop the reaction. Competent *E. coli* DH10B cells were retrieved from a -80°C freezer and transformed with 10μl of the ligated product using heat shock method (Chang *et al*., 2017). An aliquot of 100μl was spread on an LB Agar High Salt Medium plate containing 25μg/ml Zeocin and 20μg /ml X-gal and incubated at 37°C for 16h. Five blue transformed colonies were picked and inoculated in 5ml LB broth with 25μg/ml Zeocin and incubated overnight at 37°C in an orbital shaker at 250rpm. The cells from the 5 overnight cultures were harvested by centrifugation at 11000xg for 2 minutes to obtain pellet. Plasmid DNA was extracted from pelleted cells using minipreps (NucleoSpin Plasmid EasyPure Kit - Macherey-Nagel) following manufacturer’s instructions. A restriction reaction using *LguI* enzyme (Thermo Fisher Scientific) was done to confirm successful transformation with recombinant plasmid.

#### Cloning of three defined 9-LOX fragments into destination plasmid vector (pPICZαB)

The intermediary recombinant vectors *pAGM52925* containing the 548bp fragment, *pAGM52901* containing the 993bp fragment, *pAGM52914* containing the 1144bp fragment and destination plasmid vector *pPICZαB* containing the *LacZ* fragment (649bp) were used for the golden gate assembly (Figure 1c) using reaction conditions adopted from Terfrüchte *et al* (2014). Briefly, a 15μl reaction was set up containing equimolar amounts (100ng) of destination vector *pPICZαB*, and intermediary recombinant plasmid vectors *pAGM52925, pAGM52901, pAGM52914*, 3U *LguI* Type II restriction enzyme (Thermo-Fisher Scientific) that cuts through the intermediary vectors to release the 9-LOX fragments and also excises the *LacZ* fragment from the destination vector, 1X Ligation buffer (New England Biolabs), 3U of Ligase (New England Biolabs), and nuclease-free water. The reaction was performed in a thermocycler (Eppendorf Master cycler X50 Thermocycler) using the following program: 10 cycles (37°C for 2 min to allow *LguI* enzyme activity and 16°C for 5min to allow for ligase activity) and finally heat inactivation of enzymes at 80°C for 5min. The restriction-ligation product was then used to transform competent *E. coli* DH10B cells using heat shock method. Plasmid DNA extraction was done using Nucleospin plasmid Easy Kit following manufacturer’s instructions. Restriction analysis was done on the recombinant Plasmid DNA (*pAGM52943*) using *BsaI, BstXI*, and *NheI* restriction enzymes (New England Biolabs) to confirm the expected size of the insert. The plasmid DNA samples were quantified using Qubit^®^ 3.0 Fluorometer (Invitrogen) following manufacturer’s protocol. Three aliquots of the samples were sent to Eurofins Genomics, Germany for sequencing. The resultant sequences were in frame with those expected and thus one plasmid DNA sample was used for recombinant protein expression analysis.

#### Linearization of 9-LOX recombinant plasmid DNA using *Pme 1* restriction enzyme

Linearization was performed by restriction analysis using *Pme 1* enzyme in a 20μl reaction volume basing on a protocol adopted from Guo *et al* (2016). The reaction mixture contained 1X Tango buffer, 1U of restriction enzyme *Pme 1* (NEB), 50ng of 9-LOX plasmid DNA and nuclease free water. A control reaction containing unlinearized *pAGM52943* plasmid was also set up using similar reaction conditions as the test but without the *Pme 1* enzyme. The reaction mixtures for the *pAGM52943* plasmid and control were then incubated at 37°C for 3h and 5μl of the linearized and unlinearized products were checked to confirm linearization on a 1% agarose gel and viewed using a UV gel documentation system Wagtec, United Kingdom. *Pichia pastoris* X-33 strain provided as a stab in the *Pichia pastoris* expression kit (Thermo Fischer Scientific) was transformed with linearized 9-LOX plasmid DNA using electroporation method as described in the *Pichia* Easy Comp™ Transformation Kit and mixture was then incubated at 30°C without shaking for 2h to allow expression of Zeocin resistance. From the transformed cell mixture, 250μl was spread on an YPDS (Yeast Extract Peptone Dextrose) agar plate containing 25μg/ml of zeocin using sterile glass beads and left to air dry for 30min then incubated at 30°C for up to 10days. Many colonies grew but only 5 were picked using a sterile wire loop and streaked individually on five fresh YPDS agar plates containing 25μg/ml of Zeocin and incubated for 10 days at 30°C to obtain pure colonies.

#### Expression and analysis of recombinant 9-LOX protein

Protein expression was achieved by growing recombinant *Pichia pastoris* X-33 strain in Buffered Glycerol Complex Medium (BMGY) for 18h and thereafter grown in Buffered Methanol Complex Medium (BMMY) to induce protein expression. At 24h intervals, absolute methanol was added to a final concentration of 0.5% to maintain expression of recombinant 9-LOX protein. Briefly, five single colonies from the transformed *Pichia pastoris* X-33 cells were selected and each grown in 3ml of YPDS media in 50ml falcon tubes at 30°C with shaking at 250 rpm. From each of the overnight cultures, 500μl was aliquoted and inoculated into five 250ml baffled Erlenmayer flasks containing 30ml of BMGY for cell growth at 30°C with shaking at 250rpm for 18h before inducing protein expression. The cultures were then spun down at 3000xg for 5min and the pellets resuspended in 30 ml BMMY and centrifuged at 3000xg for 5min to remove excess glycerol. The resultant pellets were resuspended in 30ml of BMMY to induce protein expression and these were allowed to grow at 30°C with shaking at 250rpm for 96h. To maintain expression, 100% methanol was added to the first 4 flasks at a final concentration of 0.5% at 24h intervals for the time points 24h, 48h, 72h, and harvested at 96h. The 5^th^ flask was used as a control for uninduced expression since absolute methanol was not added to it. After expression, 30ml of the resultant culture from each of the five baffled flasks were spun at 3000xg for 5min and 1ml of supernatants pipetted off into separate Eppendorf tubes. The resultant pellet samples and the supernatants were stored at -80°C freezer until future analysis. The pellet samples were analyzed for protein expression using SDS-PAGE and Western blotting coupled with a prestained protein marker-180kDa (Thermo-Fisher Scientific).

#### SDS-PAGE analysis of recombinant 9-LOX protein

Two SDS-PAGE containing 12% and 5% separating and stacking gels respectively were prepared and poured into gel troughs to solidify. These two gels were then placed into an electrophoresis tank which was later filled with an electrophoresis buffer (Tris base 15g/l, glycine 72g/l, SDS 5g/l). The five (5) cell lysate samples (4 induced and 1 uninduced expression) were retrieved from the -80°C freezer, thawed and placed on ice. To each of 5 labelled Eppendorf tubes, 20μl of cell lysate was added followed by 20μl of the 2x SDS-PAGE gel loading buffer (4% SDS, 20% glycerol, 200mM dithiothreitol (DTT), 0.01% bromophenol blue). The contents were boiled for 10min after which 20μl from each sample was loaded in the gel wells. To the first well, 10 μl of prestained molecular weight size marker (PageRulerTM Prestained Protein Ladder, 10 to 180 kDa, Thermo Fisher Scientific) was loaded followed by uninduced sample. This was followed by 20μl of each of the four induced samples. The gels were run at 120V for 60min in an electrophoretic apparatus using tris glycine electrophoretic buffer (tris base 15g/l, glycine 72g/l, SDS 5g/l). After, one of the gels was stained with Coomassie brilliant blue (0.1% (w/v) G-250, 0.29M phosphoric acid and 16% saturated ammonium sulfate) overnight at 25°C. After, the gel was destained using destaining solution I (40% methanol and 10% acetic acid) for 30min then destaining solution II (5% methanol and 7% acetic acid) for 30min. Protein bands approximately 95KDa were observed as expected in the expressed protein sample. The second gel that was not stained was used for Western blotting (refer to gel).

#### Western blot analysis of recombinant 9-LOX protein

The separated proteins from the second SDS-PAGE gel that was not stained using Coomassie brilliant blue were electrophoretically transferred onto a solid Polyvinylidene difluoride (PVDF) membrane (Thermo-Fisher Scientific) which was later probed using anti-histidine antibody (Abcam) diluted at 1:2000 at 25°C for 1h. The PVDF membrane was then washed two times with TBS (Tris buffered saline) containing 0.05% Tween 20, each wash for 10min. The membrane was then incubated with secondary antibody (anti-mouse conjugated to alkaline phosphatase) in a 1:30000 dilution in 1x TBS containing 0.5% skimmed milk at room temperature for 1h. The membrane was then washed 3 times with 1x TBST (Tris buffered saline containing 0.05% Tween-20). Each wash was 10min at room temperature. After, the membrane was washed 2 times in alkaline phosphatase (AP) solution (0.1M Tris HCl, 0.1 M NaCl, 5mM MgCl2 and made up to pH 9.5). The membrane was stained with 15ml AP containing 20μl BCIP (Bromochloroindolyl phosphate) and 0.5% skimmed milk. Color development was monitored visually for 20min and the reaction stopped using 200μl of 1mM EDTA, pH 8.0 contained in 50ml of PBS. The expected band size of 95KDa was observed for all the 4 induced samples and was absent for the uninduced sample.

## RESULTS

### Three Amplified 9-LOX DNA fragments of *Solanum tuberosum*

The three 9-LOX DNA fragments were successfully amplified by PCR using the primer pairs which added *BsaI* sites flanking each of the three DNA fragments generated (Figure 2). The 3 fragments were individually subcloned into intermediary plasmid vector *pAGM35763*.

**Figure 2:**
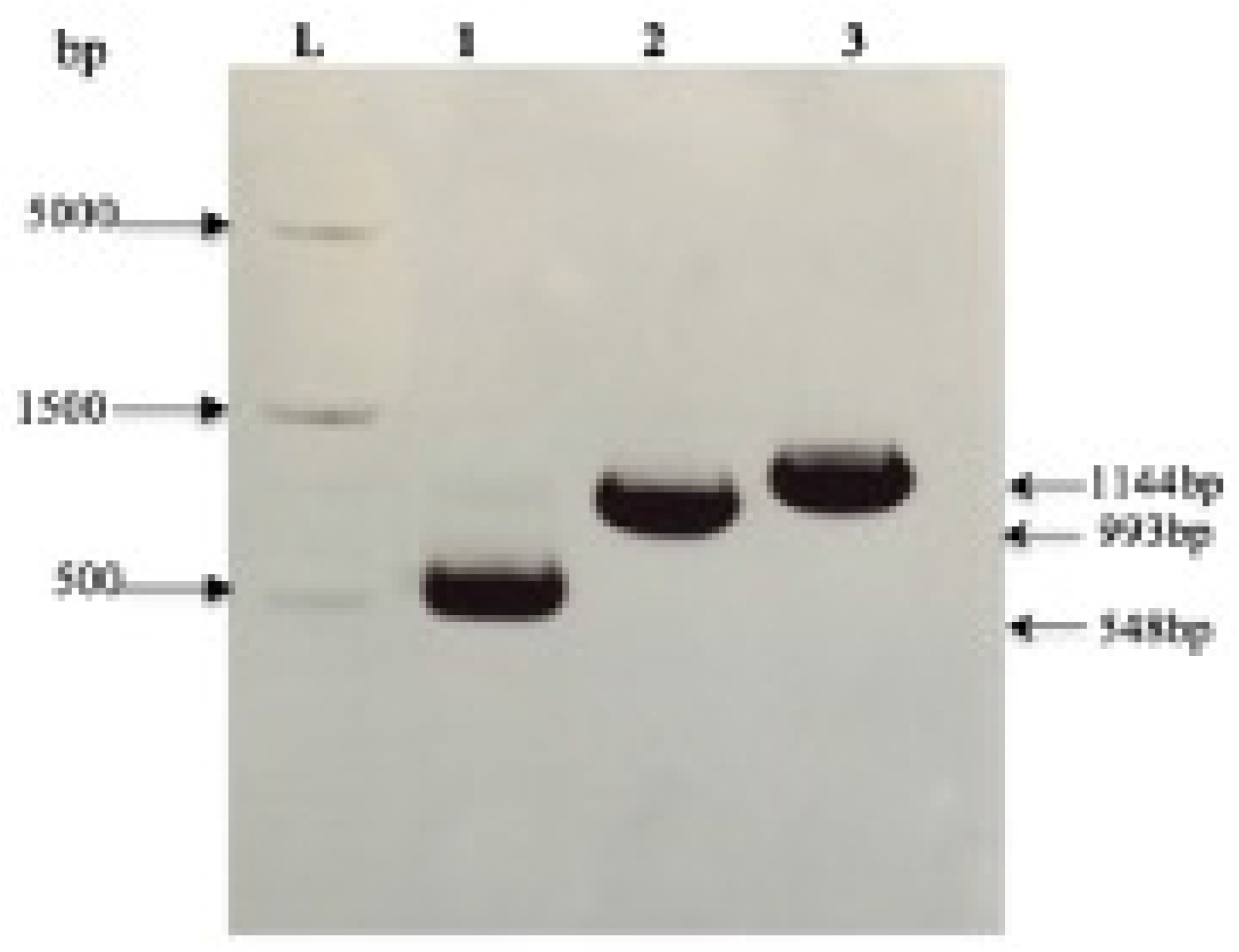
A representative 1% agarose gel showing 9-LOX amplicons. Lane L, Gene ruler 1kb plus-RTU (Thermo-Fisher Scientific), Lanes 1,2, and 3 represent the 9-LOX amplified fragments of band sizes 548bp, 993bp and 1144bp respectively. Amplifications were achieved by use of 3 pairs of primers that bind to 6 different regions of the 9-LOX DNA. After PCR amplification, 5μl of each of the three 9-LOX amplicons were mixed with 6X gel loading dye (Thermo-Fisher Scientific), loaded on a 1% agarose gel and run alongside a 1kb DNA ladder (RTU-Thermo Scientific) at 125V in 1X Tris-Acetic acid-EDTA (TAE) buffer containing 0.5μg/ml ethidium bromide for 35min. The ethidium bromide stained gel was visualized using the UV gel documentation system -Wagtec, United Kingdom.

### Amplified *LacZ* fragment from *pICH41308* plasmid vector

The *pICH41308* plasmid vector donated by Dr. Marillonnet Sylvestre was used as a template for *LacZ* gene amplification using sapla 3 forward and sapla 4 reverse primers. A 649bp amplicon was generated (Figure 3).

**Figure 3:**
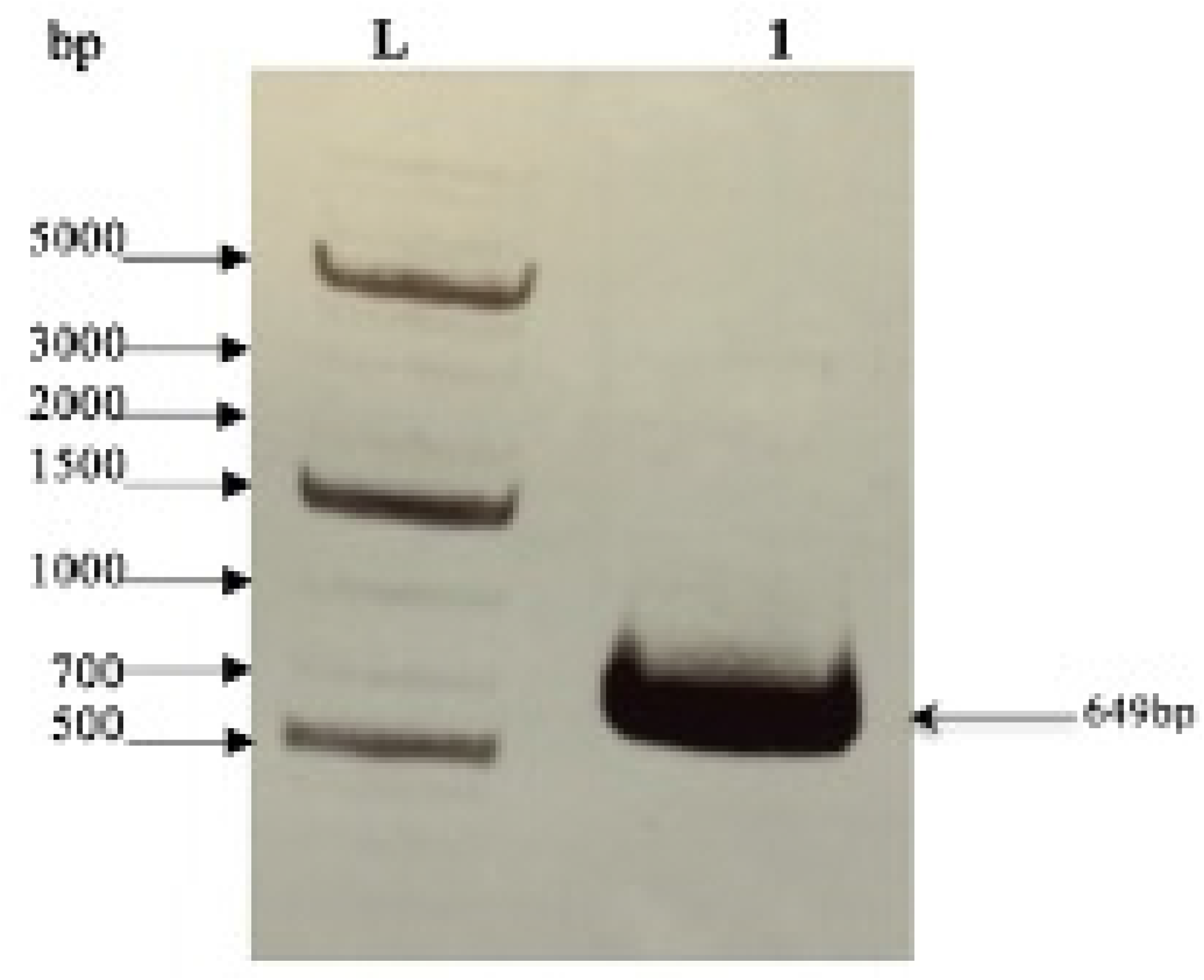
A representative 1% agarose gel showing *LacZ* amplicon. Lane L, Gene ruler 1kb plus DNA ladder RTU (Thermo-Fisher Scientific), lane 1, PCR amplicon of *LacZ* fragment size (649bp) from *pICH41308* plasmid. This amplicon was generated using PCR conditions described by Engler & Marillonnet (2013). After PCR amplification, 5μl of reaction product was mixed with 6X gel loading dye (Thermo-Fisher Scientific), loaded on a 1% agarose gel and run alongside a 1kb DNA ladder (RTU-Thermo Scientific) at 125V in 1X Tris-Acetic acid-EDTA (TAE) buffer containing 0.5μg/ml ethidium bromide for 35min. The ethidium bromide stained gel was visualized using the UV gel documentation system Wagtec, United Kingdom.

### *Pme1* linearized 9-LOX recombinant plasmid *pAGM52943*

Linearization of recombinant plasmid *pAGM52943* was achieved by using *pmel* restriction enzyme that gave a single linear band of approximately 6136bp (Figure 4).

**Figure 4:**
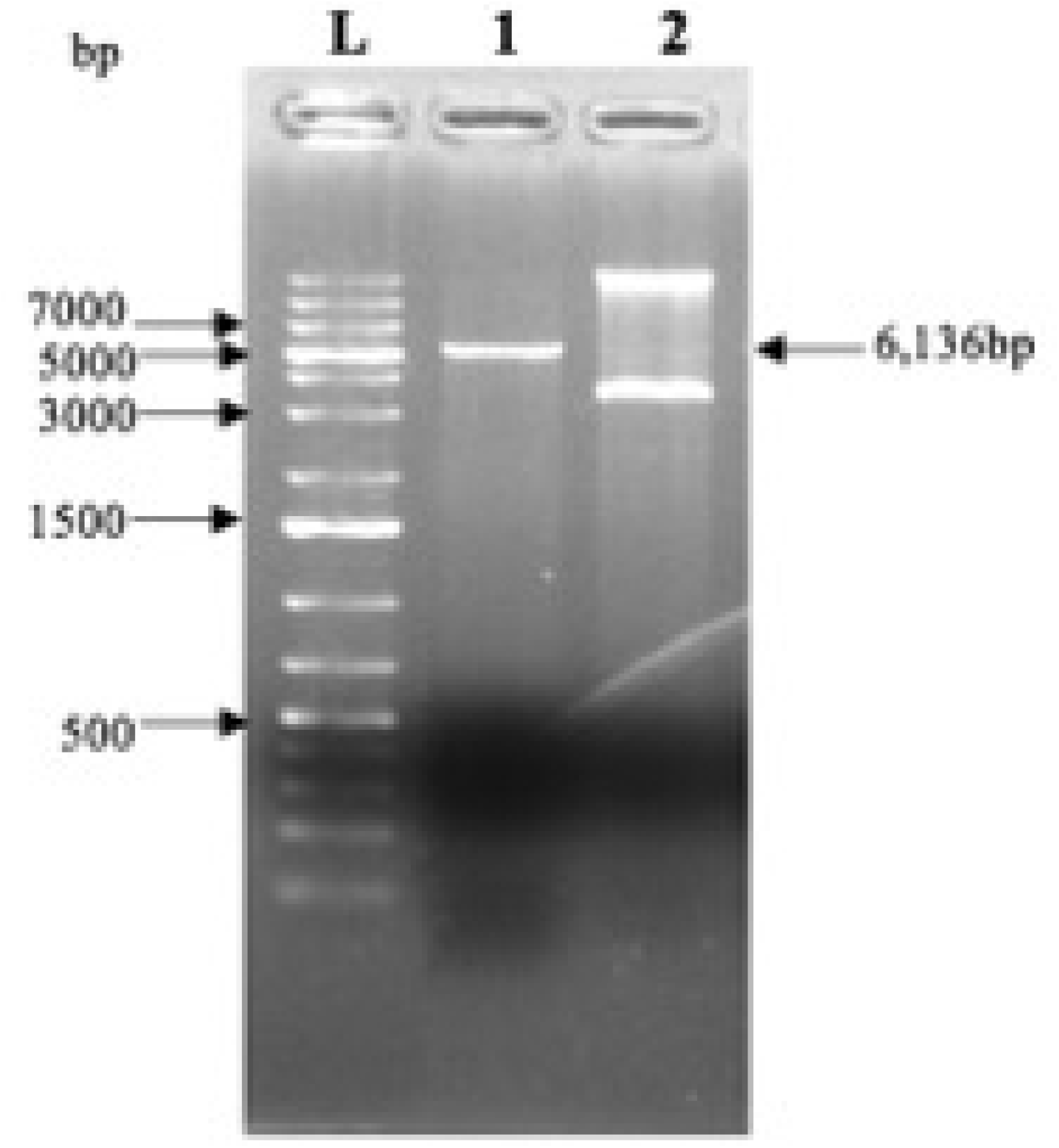
A representative 1% agarose gel with linearized and non-linearized recombinant plasmid DNA. Lane L, Gene ruler 1kb plus DNA ladder RTU (Thermo-Fisher Scientific), lanes 1 and 2 represent linearized and non-linearized *pAGM52943* plasmid vectors respectively. The reaction mixes of both linearized and non-linearized were incubated at 37°C in an incubator for 3h and after 5μl of reaction product from each tube was checked on a 1% agarose gel and viewed using a UV gel documentation system Wagtec, United Kingdom. The non-linearized plasmid was used as a control to assess linearization.

### A 12% SDS-PAGE and Western blot of recombinant 9-LOX expressed in *Pichia pastoris* X-33 strain

The recombinant 9-LOX protein has a molecular weight of approximately 97kDa. The expression of the recombinant 9-LOX protein in *Pichia pastoris* X-33 cells was analyzed by SDS-PAGE and Western blotting and showed bands from cell lysate samples (Figure 5).

**Figure 5:**
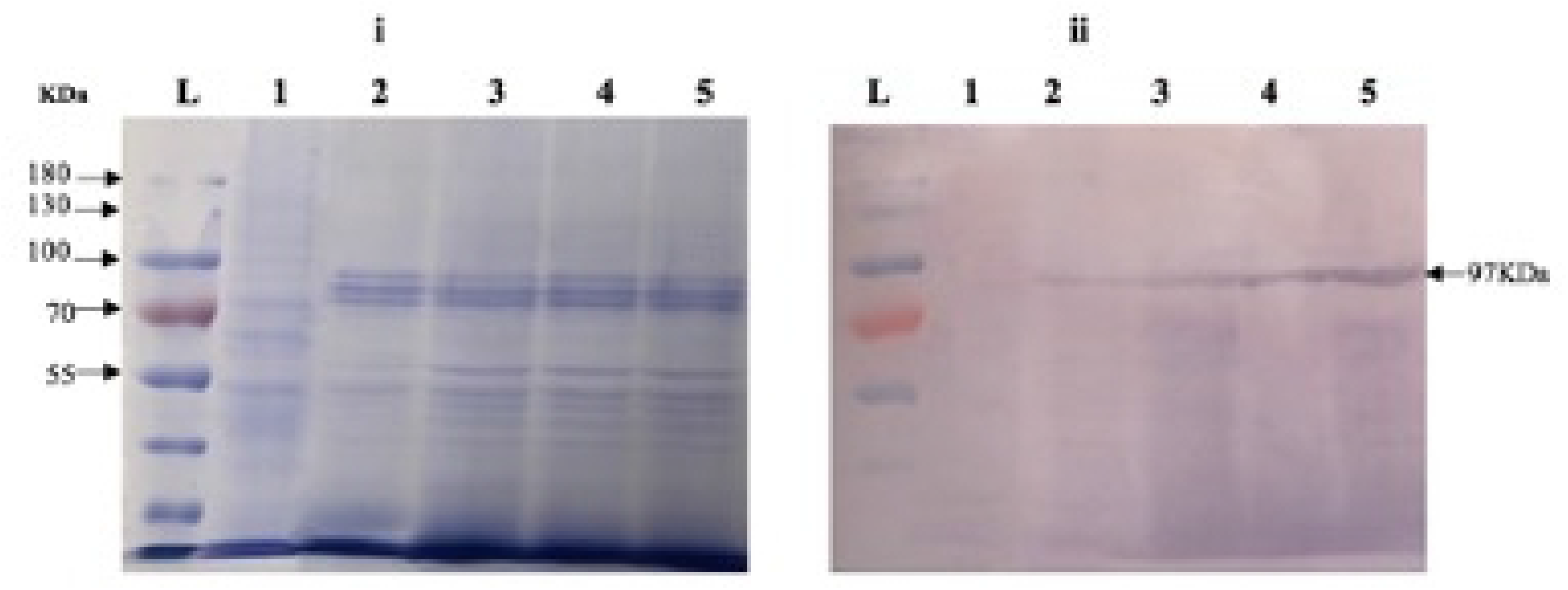
A 12% SDS-PAGE and Western blot of recombinant 9-LOX expressed in *Pichia pastoris*. A Coomassie blue staining of polyacrylamide gel (i): Lane L, Prestained protein marker-180kDa (Thermo-Fisher Scientific), lane 1, uninduced recombinant *Pichia pastoris* cells at 96h post expression. Lanes 2, 3, 4, and 5 represent proteins from induced *Pichia pastoris* cells integrated with *pAGM52943* plasmid at 96h post induction with 100% methanol to a final concentration of 0.5%. All the four induced samples showed expected band size of 97KDa. Western blot analysis of recombinant 9-LOX protein gel (ii). Lane L, Prestained protein marker-180kDa (Thermo-Fisher Scientific), lane 1, PVDF (Thermo-Fisher Scientific) transferred proteins from uninduced *Pichia pastoris* cells integrated with *pAGM52943* plasmid at 96h of expression. Lanes 2, 3, 4 and 5 represent proteins from induced cells containing *pAGM52943* plasmid at 96h post induction with 100% methanol to a final concentration of 0.5%. Western transfer probed with anti-His monoclonal antibodies resulted in bands of 97KDa that correspond to the recombinant 9-LOX protein. Non induced sample showed no signal.

## DISCUSSION

This study aimed at generating recombinant *Solanum tuberosum* 9-LOX protein using *Pichia pastoris* as an expression system. For many years, plant pathogenic fungi have caused devastating losses worth millions of dollars in crops worldwide (Yoon *et al*., 2013). Synthetic fungicides have for many years controlled plant pathogenic fungi. However, their repeated use over decades has disrupted natural biological systems, and sometimes resulted in development of fungal resistance (Yoon *et al*., 2013). They have undesirable effects on non-target organisms, and foster environmental and human health concerns and thus alternative approaches that yield biodegradable fungicides have to be investigated. This study has successfully proven that *Pichia pastoris* can generate recombinant 9-LOX protein and recombinant cell lines can be prepared for production of large quantities of recombinant 9-LOX protein. Golden Gate cloning was used due to the large size of the 9-LOX insert (2629bp) that codes for a protein of approximately 97kDa. The vector NTI Bioinformatics tool Version 11.5.5 was used to select the three 9-LOX fragments that were subcloned into intermediary level 0 vector as described by Engler & Marillonnet (2013). Assembly of the 9-LOX PCR products directly in the destination vector without first cloning them into intermediary vectors is possible but it is recommended to purify the PCR products using a column to remove any DNA polymerase left in the PCR product and to remove primer dimers that may be produced during PCR amplification. Some of the primer dimers can be flanked by two fusion sites (these are part of the primers) and can thus be cloned, which will always result in incorrect constructs (Werner, Engler, Weber, Gruetzner, & Marillonnet, 2012). This study was able to clone a 2629bp 9-LOX gene into pPICZαB plasmid vector using restriction-ligation reaction for assembly of three 9-LOX fragments into the destination plasmid vector. The golden gate reaction conditions used in this study are similar to those described by Marillonnet and Grützner (2020) and Terfrüchte *et al* (2014).

The level 0 intermediary vector *pAGM35763* was cloned with individual 9-LOX fragments of 548bp size attached to a 6xHistag, 9-LOX fragment of 993bp, and 9-LOX fragment of 1144bp. All the 3 fragments cloned into level 0 intermediary vectors carried BsaI restriction sites introduced during PCR amplification. The level 0 intermediary vector carrying each of the three 9-LOX fragments was then used in a final restriction-ligation step in which the fragments were assembled into a level 1 destination vector (pPICZαB) which generated recombinant plasmid *pAGM52943*. All level 0 intermediary vectors met the following criteria as described by Engler & Marillonnet (2013): (1) They were flanked with *BsaI* sites in opposing direction. The enzyme *BsaI* (isoschizomer: Eco31I) with the non-palindromic recognition/hydrolysis site is active at both 37°C and 50°C, has a recognition site of 6 bp separated by one random nucleotide from the site of hydrolysis and generates a 4 bases overhang (5’ -GGTCTC(N)1-3’ / 3’-CCAGAG(N)5-5’, (2) all DNA parts of the same type were flanked by same pair of fusion sites, (3) recognition sequences of the restriction endonuclease *BsaI* were absent in all internal sequences of the selected 9-LOX fragments, (4) the pPICZαB vector backbone contained a *LacZ* resistance gene for negative selection introduced by T4 polymerase and ligase buffer without using ligase. These guiding cloning principles were also similar to those described by Marillonnet and Grützner (2020). According to Jeong *et al* (2012), T4 cloning was used to undertake a one-step sequence and ligation independent cloning similar to this study since ligase enzyme was not used in the cloning reaction of the Lac Z gene into the digested pPICZαB plasmid vector. This is attributed to the fact that T4 DNA polymerase has a 3’ to 5’ exonuclease activity (Jeong *et al*., 2012). Sequencing of the recombinant destination plasmid vector *pAGM52943* showed no mutations in the shuffled 9-LOX gene fragments. This was expected because the modules in the intermediary vector constructs were made using PCR but then sequenced before being used for restriction-ligation assembly and correct orientation was noted.

The recombinant *pAGM52943* plasmid was used for protein expression using the methylotrophic *Pichia pastoris* X-33 strain as a host system. The recombinant 9-LOX protein was expected to be secreted in the supernatant since the 9-LOX gene was cloned upstream of the alpha secretion factor using the golden gate protocol as described by Engler & Marillonnet (2013). Attempts to recover recombinant 9-LOX protein from supernatant were futile which could be attributed to vacuolar proteases such as carboxypeptidase Y and proteinase B which are significant factors in protein degradation especially in fermenter cultures, owing to the high cell density environment, in combination with lysis of small percentage of cells (Cregg *et al*., 1989). An example of a strategy to counter proteolytic degradation of secreted recombinant proteins is to use mutant strains deleted in the vacuolar proteases saccharopepsin (Protease A) and/or cerevisin (Protease B). These proteases are mainly present in the extracellular medium as a result of cell lysis during protein expression observed by frothing in the baffled flasks (Werten *et al*., 2019). Methanol can also induce cell lysis by oxidative stress and heat-shock responses, eliciting a proteolytic response when the cells are growing exponentially, which results in high cell-density fermentation (Vieira *et al*., 2017). According to Vieira *et al* (2017), the mass spectrometry analysis of protein expression in the medium identified serine proteases and intracellular proteins of yeast in addition to recombinant rDM64 protein which suggested that yeast lysis occurred during expression in shake flasks. Parameters such as starvation, heat, pH changes, and toxic chemicals affect the growth of *Pichia pastoris* in shake flasks. In their expression study therefore, Pefabloc, a potent and irreversible serine proteinase inhibitor was added to the expression medium. This serine protease inhibitor was used to protect against the proteolytic degradation induced by yeast serine proteases although it did not completely eliminate the protease activity.

In this study, the 9-LOX recombinant protein was recovered from the cell lysates. SDS-PAGE analysis showed a band size of approximately 97KDa of expected band size that was confirmed by immunoblotting of PVDF membrane probed using anti-histidine antibody which is very good for lowly expressed proteins although it gives a high background (Xiang *et al*., 2021). The expression results from this study are similar to those from a study by Rodrigo *et al* (2013) where expressed recombinant protein was recovered from the cell lysates as was revealed by SDS-PAGE and Western blot analysis. During decades of using *Pichia pastoris* as a eukaryotic protein expression system, the problem of inconsistent secretory productivity among different recombinant proteins, that is to say some proteins could reach extremely high yields while some others had little or no expression at all. This has always been a major obstacle for routine application in both research and industry *(Yang et al*., 2013). The same study by Yang *et al* (2013) suggested a new strategy to increase *P. pastoris* secretory productivity by optimizing the yeast convertase Kex2 cleavage which increased secretory protein in comparison with yeast vector with no Kex2 modification.

Yang *et al* (2013) replaced the glutamic acid at the kex2 cleavage site with each of the other 19 natural amino acids in plasmid vector pPICZaA, and the effect on protein expression was assessed using luciferase as a reporter gene. All the recombinant plasmids in their study were transformed into the X-33 strain. Single copy strains were selected based on their Zeocin resistance ability, and clones were grown and induced with methanol. Measurement of fluorescence by the plate reader showed that replacement of Glu with Ser gave 13-fold higher expression than the mutant with Tyr. Also, artificial addition of the Kex2 protease gene into the host genomic DNA expressing Venus fluorescence protein doubles the productivity of the recombinant protein according to a study by Juturu & Wu (2018).

In this study, expression of recombinant *Solanum tuberosum* 9-LOX protein from cell lysates was achieved using *Pichia pastoris* X-33 strain. Therefore, to enable secretion of recombinant 9-LOX protein from *Pichia pastoris*, optimization of the α-secretion factor and yeast kex2 convertase in comparison to yeast vector with no kex2 modification can be investigated. Artificial addition of the Kex2 protease gene into the host genomic DNA should be investigated during expression to enhance recombinant protein secretion into the culture media and protease inhibitors should be included during expression to avoid loss of protein due to proteolytic activity of proteases in the culture media. Potential testing of the recombinant 9-LOX protein for its chemoenzymatic activity should also be done.

## Acknowledgments

Funding was provided by the German Federal Ministry of Education and Research under the OXiLiFungi Project: 13FH0261×5. Extra funding was provided by DAAD Student Academic Exchange Program. Thanks to the Leibniz Institute of Plant biochemistry for providing lab space for performing the cloning work. Thanks to the Molecular Biology Lab of College of Veterinary Medicine, Animal Resources and Biosecurity, Makerere University for providing Lab space and equipment for conducting protein expression studies under the anti-tick Vaccine Project.

## Authors’ contribution

Conceptualization, funding, and project administration were done by Professor Leif-Alexander Garbe, Dr. Fabien Schultz, and Matthias Koch. Methodology and investigation were done by Alex Olengo, Ann Nanteza, Elvio Henrique Benatto Perino, Sylvestre Marillonnet, and Ramona Grützner. Cloning constructs were provided by, Dr. Marillonnet Sylvestre, Dr. Ramona Grützner and Elvio Henrique Benatto Perino. Project supervision was done by Dr. Ann Nanteza and Professor Leif-Alexander Garbe. Dr. Margaret Saimo-Kahwa provided protein work consumables. Original draft was written by Alex Olengo and Ann Nanteza. Reviewing and editing was performed by Alex Olengo, Ann Nanteza, Elvio Henrique Benatto Perino, and Dr. Saimo-Kahwa.

## Competing interests

The Authors declare that they have no competing financial interests or personal relationships that could have appeared to influence the work reported in this paper.

## Notes

### Competing Interest Statement

The authors have declared no competing interest.

